# Abdominal pain assessment in rabbits: using the CANCRS to recognize pain and testing its internal validity over time

**DOI:** 10.1101/2020.10.27.356832

**Authors:** Penelope Banchi, Giuseppe Quaranta, Alessandro Ricci, Mitzy Mauthe von Degerfeld

## Abstract

A composite scale for pain assessment in rabbits has been previously designed and tested (CANCRS). The present study describes the refinement of the scale and the evaluation of its ability to detect pain variations over time. Furthermore, a comparison between the CANCRS and the Visual Analogue Scale (VAS) has been performed, to underline the differences between an objective (CANCRS) and a subjective (VAS) assessment of abdominal pain. In the first part of the study, 86 rabbits (n=47 heathy patients and n=39 patients with gastrointestinal stasis syndrome) underwent pain assessments with the VAS and the CANCRS. Thirty-two patients with gastrointestinal stasis syndrome participated to the second part of the study. These patients underwent four pain assessments with the CANCRS. The first assessment took place before meloxicam administration and the others after 30, 60 and 90 minutes. The CANCRS showed differences between healthy and diseased rabbits (*P* = 0.0001), median scores were 5 (IQR 4 - 6) and 9 (IQR 7 - 11) respectively. The VAS showed differences between healthy and diseased rabbits (*P* = 0.02), the median scores were 4 (IQR 2 - 5.35) and 5.3 (IQR 2.65 - 6.45) respectively. The cut-off scores for the CANCRS and for the VAS for differentiation between healthy and diseased patients were 7 (Sp 89%, Se 79%) and 4.4 (Sp 59%, Se 69%) respectively. Sensitivity and specificity for each parameter of the CANCRS were calculated, in order to obtain weighting factors. Accordingly, the evaluation of respiratory pattern and vocalizations should be excluded from the CANCRS, since their performances in pain evaluation are poor. Internal validity of the CANCRS was tested assessing pain before and after the analgesic treatment and the results showed significancy at each time point. The CANCRS showed better performances than the VAS and its responsiveness to pain variations has been verified.

## Introduction

As rabbit popularity as companion animal increases, veterinarians’ confidence in revealing rabbit pain has improved in the last decade (1). However, most of veterinarians (64%) declare to be just “Fairly confident” and 29% declare to be “Not very confident” in revealing rabbit pain (2). A survey by Benato et al. (2020) showed how the majority of respondents relies on behavioral changes when assessing pain. On the contrary, changes in physiological parameters are considered less reliable as they can be altered in any stressful situation (3). Behavioral and physiological parameters can be included in composite pain scales (4-7), in order to point out the multidimensional aspect of pain. It has been previously showed that composite pain scales are more reliable than simple one-dimensional pain scales like the Visual Analogue Scale (VAS) (8). The VAS is a subjective tool (9) and the rater is asked to make an estimate of the pain of the patient placing a mark on a 10 cm-line. The VAS is an effective tool in human medicine (10), when a self-assessment is performed, whereas its use in veterinary medicine comes with limitations, since the assessment is performed on another individual belonging to another species. Therefore, one-dimensional scales are considered suboptimal tools for pain evaluation in veterinary medicine (11). Nevertheless, they are sometimes used in domestic animals (8, 12), but their performances have never been previously evaluated in rabbits. Testing a subjective tool like the VAS could be useful to point out the advantages and the limitations of a subjective assessment of pain. Finding a tool with high validity, reliability and feasibility is a key point in the pain management of the rabbit, in order to support the veterinarian in clinical decisions concerning analgesia and in order to monitor the patient during the hospitalization. In our previous study (4) a composite pain scale for the assessment of pain in pet rabbits was developed and its construct validity and reliability were assessed. However, internal validity should be assessed, and a refinement of the scale should be performed in order to have a useful tool (13).

One of the most common presentations for rabbits is gastrointestinal stasis (RGIS) (14, 15). It is a syndrome characterized by decreased appetite or anorexia, decreased or absent fecal output, abdominal discomfort and lethargy (15, 16). It has a multifactorial etiology (17) and it can result from dental diseases, dietary changes, stressful events, environmental changes, dysbiosis, pain, infection, neoplasia, drug effect (anesthetic agents, anticholinergics, opioids, and antibiotics), obstruction. The reduced food intake leads to retarded colonic transit, dehydration of the *digesta* and impairment of the enteric microflora. If the patient is left untreated, this process can lead to obstruction and accumulation of gas within the gastrointestinal tract, leading to severe consequences, such as dyspnea and hypovolemic shock (15).

Managing pain during gastrointestinal stasis syndrome is essential to promote food intake in rabbits. However, analgesia can have risks associated with the chosen drug. The use of opioids is discouraged due to their effects on gastrointestinal motility, that is furtherly reduced (3, 18). Non-steroidal anti-inflammatory drugs (NSAIDs), such as meloxicam, show adverse effects in dogs and cats and should be cautiously administered in rabbits because patients presenting with gastrointestinal stasis syndrome are likely to have or to develop gastric ulcerations (15, 17). A dosage of 1 mg/kg is considered safe for rabbits when administered orally with no clinical, biochemical or postmortem abnormalities reported (19). When a gastrointestinal syndrome is ongoing, oral therapy is discouraged in some cases (i.e. obstruction, altered mentation), and all the drugs should be administered parenterally (15).

Being gastrointestinal stasis is a common syndrome causing abdominal pain in domestic rabbits (16), patients presenting with the syndrome were the chosen subject of the present study. Furthermore, the majority of the respondents to the survey of Benato et al. (2020) declares to make subjective assessments for pain in rabbits. This led to the decision of testing the VAS, which is a tool that allows a subjective assessment of pain, with numerical results. Specifically, the objectives were:

I. To define cut-off scores for the CANCRS and the VAS between healthy animals and rabbits experiencing pain and to estimate their sensibility and specificity in pain detection.
II. To evaluate each parameter included in the CANCRS for its sensibility and specificity in detecting pain, for future refinement of the scale by attributing a lower weight to parameters that lacks in specificity or sensibility.
III. To assess the responsiveness of the CANCRS to changes in pain levels associate with the administration of the analgesic drug (meloxicam 1 mg/kg SQ q24) by evaluating the internal validity of the CANCRS.

## Materials and methods

Eighty-six rabbits admitted to the C.A.N.C. (the exotics and wild animals Veterinary Teaching Hospital of the Veterinary Science DPT. Of University of Torino) were enrolled in the study, between January and August 2020. No restrictions were placed on breed, sex, age or weight of the rabbits. Thirty-nine patients were admitted due to gastrointestinal stasis syndrome (RGIS), whereas 47 patients were admitted for checkups or vaccination and were deemed as healthy according to the clinical examination (HEALTY).

The owners of 7 of the 39 patients belonging to the RGIS group refused hospitalization. Therefore, only 32 rabbits of the RGIS group were included in the second part of the study. Furthermore, rabbits requiring nasogastric tube were excluded from the study because of its interference with the use of facial expression for pain evaluation.

The first part of the study took place at admission: all the rabbits underwent a pain assessment with both the CANCRS and the VAS. Regardless of the attending clinician, the same veterinarian performed all the assessments using the VAS and the CANCRS at the end of the clinical examination. The VAS score was recorded prior to the assessment of the patients with the CANCRS, in order to avoid bias deriving from clinical data collection.

Pain assessments for the first part of the study always happened in the clinical examination room. The patients were taken out form the carrier and positioned on a table for clinical examination and pain assessment.

The second part of the study included thirty-two of the patients included in the RGIS group. These patients underwent four pain evaluation with the CANCRS during the hospitalization. A pain assessment was performed just before the administration of the anti-inflammatory drug (meloxicam 1 mg/kg SQ q24). Afterwards, three assessment were performed 30 (T30), 60 (T60), and 90 (T90) minutes after the administration of the anti-inflammatory drug. Pain assessments for the second part of the study always happened in the hospitalization room. The patients were taken out from the cage and positioned on a table for clinical examination and pain assessment.

Environmental conditions, including light, noise, and temperature were stable through the process.

### Pain assessment tools

The VAS consisted of a 10-cm line representing no pain at the left end and severe pain at the right end. The rater placed a mark on the line that corresponded to the degree of pain for the patient (20, 21).

As for what concern the CANCRS, the scores were given as stated in our previous study (4) and the scale is presented in *Table 1*. It includes five facial action units (orbital tightening, cheek flattening, nostril shape, whisker position, and ear position) and each one was scored according to whether it was not present (score 0), moderately present (score 1) and obviously present (score 2).

**Table 1.**
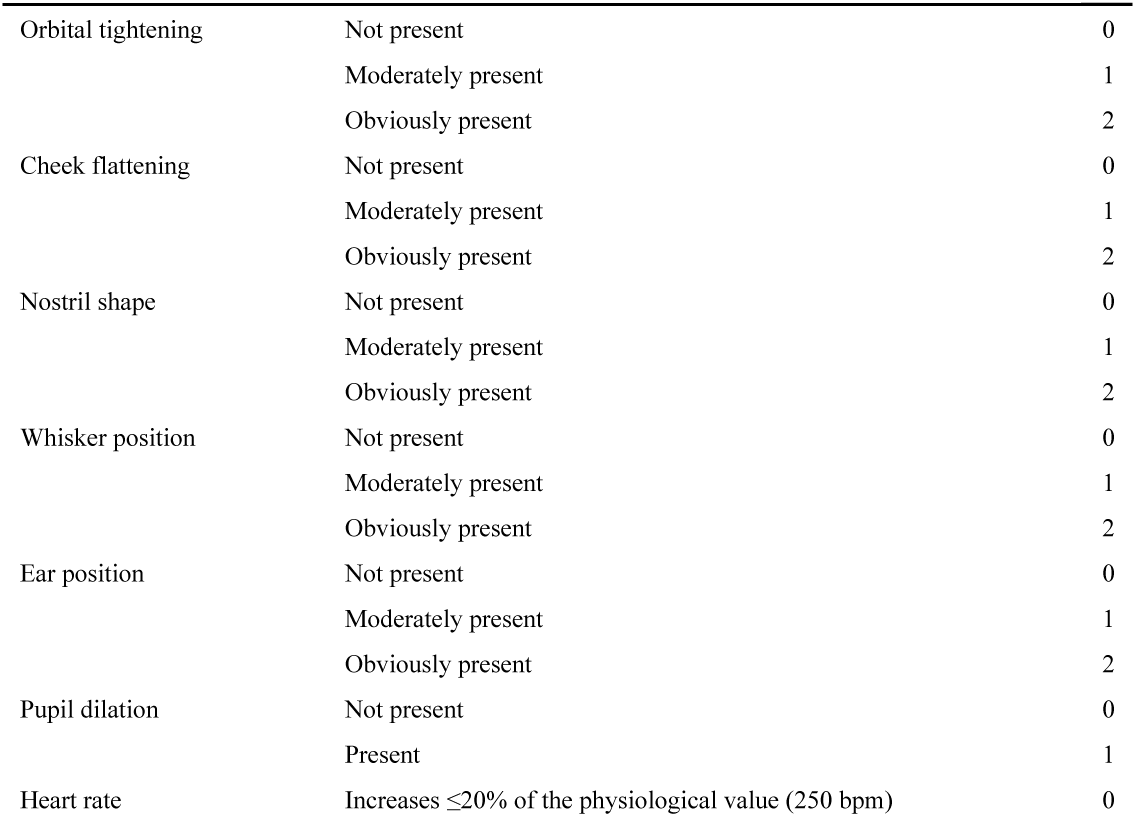
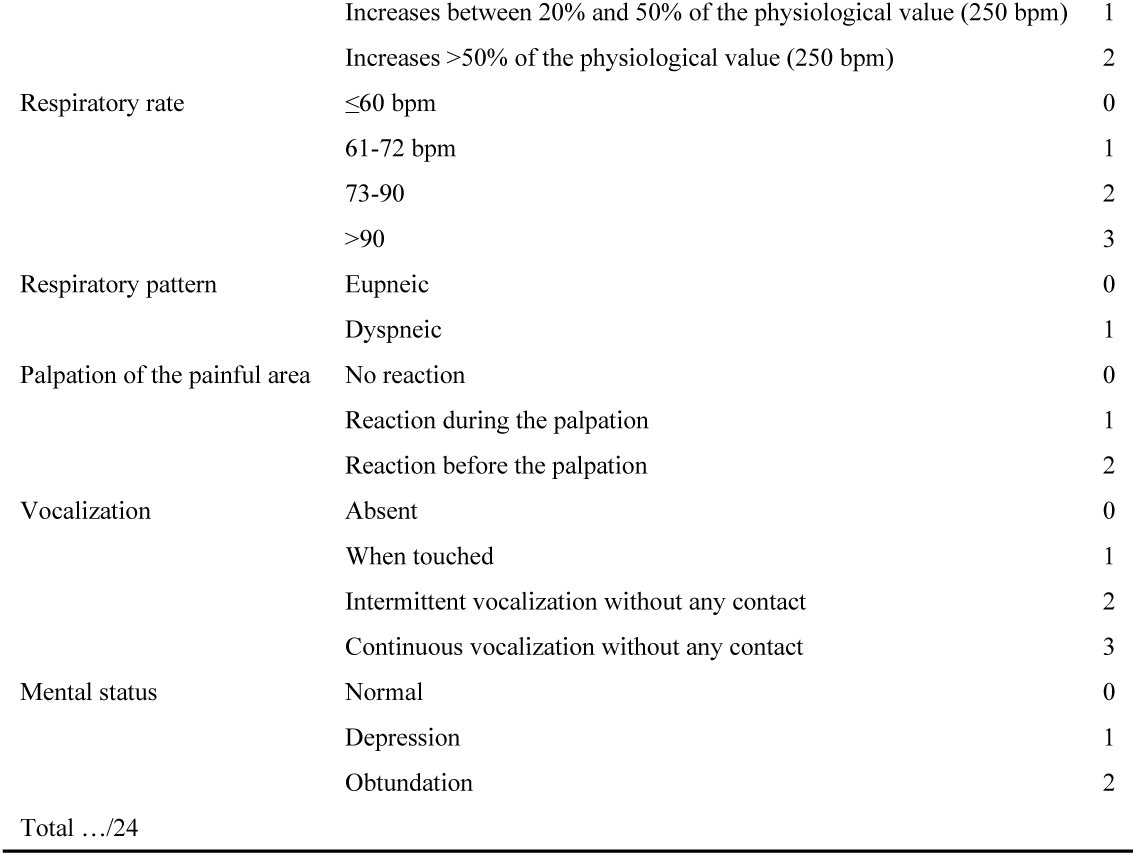
Score sheet for the CANCRS (Centro Animali Non Convenzionali Rabbit Scale).

Furthermore, it includes some clinical parameters. Pupillary dilation was scored as 1 if present or scored as 0 if not present. For heart rate, increases lower than 20% of the physiological value (250 bpm)(16) were given the score of 0, increases higher than 20% were given the score of 1, and increases higher than 50% were given the score of 2. Heart rate was measured with a stethoscope for 15 seconds and the result was multiplied by four, in order to obtain beats per minute.

The average respiratory rate varies from 30 to 60 bpm (16); for rates of 60 bpm or less a score 0 was given, for rates between 61 and 72 a score 1 was given, for rates between 73 and 90 a score of 2 was given, and for rates higher than 90 bpm a score of 3 was given.

Respiratory pattern was considered as eupneic (score 0) or dyspneic (score 1).

Since the palpation of the suspected painful area is a parameter included in the CANCRS, abdominal palpation was delicately performed and the reaction of the patient was classified as no reaction (score 0), reaction during the palpation (score 1) and reaction before the palpation (score 2).

Absence of vocalization was scored as 0, vocalization when touched was scored as 1, intermittent vocalization without any contact with the operator was scored as 2, whereas continuous vocalization was scored as 3.

Finally, the mental status was classified as ‘normal’ (score 0), ‘depression’ (score 1) and ‘obtundation’ (score 2).

In accordance with the Dipartimento di Scienze Veterinarie of Università degli Studi di Torino policy, this research does not require the approval of the ethical committee, for our assessment tool does not interfere with the clinical development of the included cases and rabbits were manipulated for routine diagnostic purposes only. Proper informed consent was collected from the owner prior to each clinical examination.

### Statistical analysis

Differences in scores between heathy animals and rabbits with RGIS were analyzed using the Mann–Whitney *U* test. Cut-off values for CANCRS and VAS were determined to obtain maximal differentiation between HEALTHY and RGIS. The same cut-off values were used to determine internal sensitivity, specificity, and positive and negative predictive values for both CANCRS and VAS. Sensitivity and specificity for individual parameters of the CANCRS were also determined. Based on these values, weighting factors for future adjustments of the scale, were determined retrospectively. When sensitivity or specificity was ≤25%, a weighting factor of 0 was applied; between 25% and 50% the weighting factor was 1; between 50% and 75% the weighting factor was 2; and when both sensitivity and specificity were ≥75% a weighting factor of 3 was applied. The effect of the analgesic drug over time for the CANCRS in RGIS patients was assessed by means of the repeated measures ANOVA followed by Bonferroni’s post hoc test.

Statistical analysis was performed using R 3.2.2. Statistical significance was accepted for *P* ≤ 0.05.

## Results

### Differences between healthy patients and rabbits with RGIS

The CANCRS showed extremely significant differences between HEALTY and RGIS rabbits (*P* = 0.0001), the median score was 5 for HEALTHY rabbits (IQR 4 to 6) and 9 for RGIS rabbits (IQR 7 to 11).

The VAS showed statistically significant differences between HEALTHY and RGIS rabbits (*P* = 0.02), the median score was 4 (IQR 2 to 5.35) and 5.3 for RGIS rabbits (IQR 2.65 to 6.45).

### Cut-off values between healthy patients and rabbits with RGIS

The cut-off value for the CANCRS for differentiation between HEALTHY and RGIS rabbits was 7. Sensitivity and specificity were 89% and 79% respectively. The AUC was 0.88 (CI 0.80-0.97). Positive predictive value was 86% and negative predictive value was 84% with accuracy of 85%. The cut-off value for the VAS for differentiation between HEALTHY and RGIS rabbits was 4.4. Sensitivity and specificity were 69% and 59% respectively, as represented in *Figure 1*. The AUC was 0.64 (CI 0.52-0.76). Positive predictive value was 49% and negative predictive value was 61% with accuracy of 53.5%.

**Fig 1.**
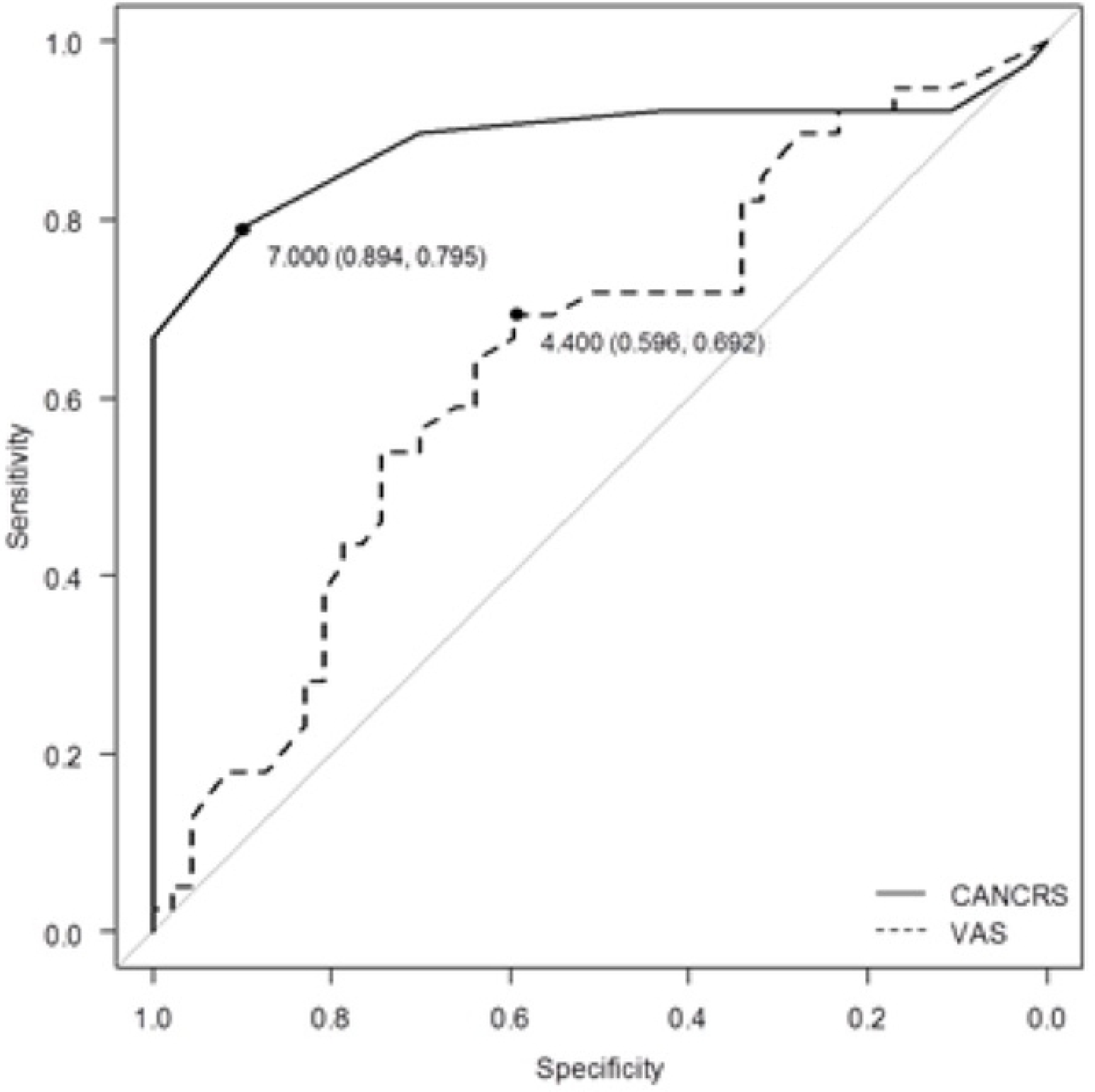
Receiving operating characteristic (ROC) curve assessing the cut‐off point for the CANCRS and the VAS to evaluate the presence of abdominal pain in rabbits. The cut-off score for differentiation between healthy rabbits and patients with gastrointestinal stasis syndrome is 7 (sensitivity 89%, specificity 79%). The cut-off score for differentiation between healthy rabbits and patients with gastrointestinal stasis syndrome is 4.4 (sensitivity 69%, specificity 59%).

Cut-off values for individual parameters of the CANCRS for differentiation between HEALTHY and RGIS rabbits are resumed in *Table 2*.

**Table 2.** Sensitivity, specificity, area under the curve (AUC) with confidence intervals (CI), cut-off values and weighting factors for individual parameters of the CANCRS.

| Parameters | HEALTHY - RGIS |  | Cut-off values | AUC | CI | Weighting factor <sup>a</sup> |
| --- | --- | --- | --- | --- | --- | --- |
|  | Sensitivity | Specificity |  |  |  |  |
| Orbital tightening | 66% | 85% | 1 | 0.77 | 0.68-0.86 | 2 |
| Cheek flattening | 56% | 80% | 1 | 0.7 | 0.60-0.80 | 2 |
| Nostril shape | 82% | 66% | 1 | 0.77 | 0.68-0.86 | 2 |
| Whisker position | 77% | 76% | 1 | 0.79 | 0.7-0.87 | 2 |
| Ear position | 51% | 57% | 1 | 0.53 | 0.41-0.64 | 2 |
| Pupil dilation | 77% | 68% | 1 | 0.72 | 0.63-0.82 | 2 |
| Heart rate | 64% | 70% | 2 | 0.66 | 0.55-0.77 | 2 |
| Respiratory rate | 100% | 61% | 2 | 0.76 | 0.66-0.86 | 2 |
| Respiratory pattern | 21% | 98% | 1 | 0.59 | 0.52-0.66 | 0 |
| Palpation of the painful area | 28% | 91% | 1 | 0.6 | 0.51-0.68 | 1 |
| Vocalization | 7% | 100% | 1 | 0.49 | 0.49-0.58 | 0 |
| Mental status | 53% | 91% | 1 | 0.73 | 0.64-0.82 | 2 |

<sup>a</sup>Sensitivity or specificity is $\leq 25\%$ , weighting factor = 0; 25-50%, weighting factor = 1; 50-75%, weighting factor = 2; sensitivity and specificity are $\geq 75\%$ , weighting factor = 3.

### Internal validity of the CANCRS

Patients experienced a strongly significant reduction in pain at each time point following meloxicam administration (P≤0.001 between T0-T30, T0-T60, T0-T90, T30-T60, T30-T90, T60-T90). Means (±DS) of the CANCRS scores for each time point are resumed in *Table 3* and the variation of pain scores over time is presented in *Figure 2*.

**Table 3.** Means and standard deviations of the CANCRS scores at each time point. Assessments were performed just before the meloxicam administration (T0) and 30 (T30), 60 (T60), 90 (T90), and 120 (T120) minutes after the administration.

|  | Mean | DS |
| --- | --- | --- |
| T0 | 9.37 | 1.71 |
| T30 | 8.43 | 1.99 |
| T60 | 7.12 | 2.07 |
| T90 | 5.18 | 1.82 |

**Fig 2.**
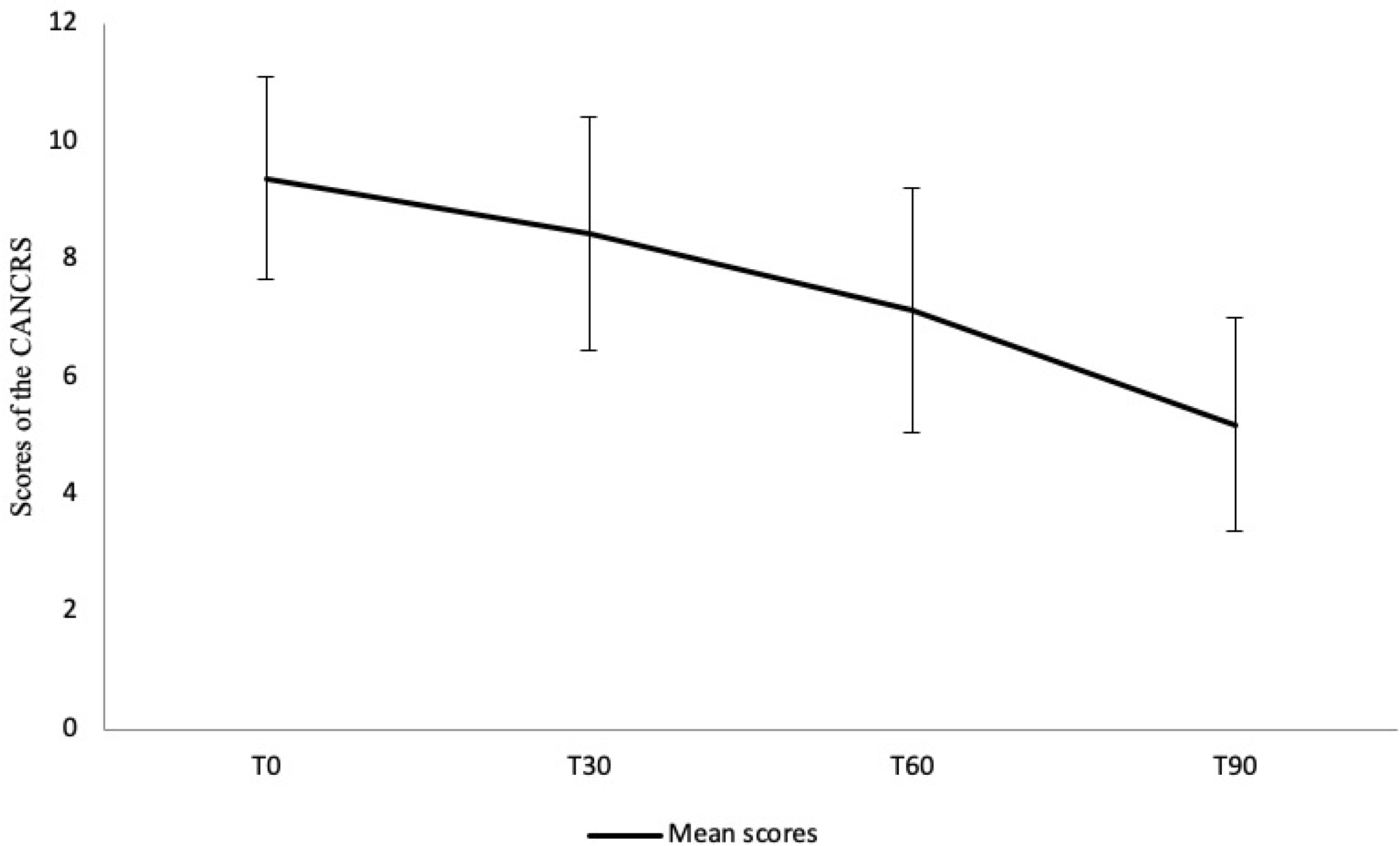
Variation of CANCRS pain scores over time. The mean scores (±DS) of the scores of the CANCRS from T0 to T90 is represented in the figure. It can be observed that patients experienced a significant reduction in pain at each time point following meloxicam administration (P<0.05 between T0-T30, T0-T60, T0-T90, T30-T60, T30-T90, T60-T90).

## Discussion

The present study describes the use of the CANCRS and the use of the VAS to detect pain in rabbits with gastrointestinal stasis syndrome, the responsiveness of the CANCRS to analgesic treatment, and it takes another step towards the refinement of the scale. Difference in pain levels between healthy rabbits and patients with abdominal pain can be detected with both tools. The CANCRS scale shows a cut-off score of 7 to differentiate between healthy animals and rabbits with abdominal pain with good sensitivity and specificity and it can detect pain variation over time according to analgesia administration. Even if pain variation over time were not investigated for the VAS, the scale showed very low sensibility and specificity in rabbits. No previous study concerning the VAS in rabbits had been performed previously, however the limitations of the VAS were previously discussed for other veterinary species (8, 12, 22-24). The VAS is a subjective, simple and poorly reliable tool and it is not aimed to consider the multidimensional aspect of pain (25), whereas composite pain scales are objective tools and are intended to capture the behavioral and physiological changes associated with pain. According to Benato et al. (2020), the majority of veterinary surgeons is fairly confident in recognizing pain in rabbits, although it is mostly a subjective evaluation. This led to the decision to test the VAS for pain detection and quantification in rabbits, for it allows a subjective assessment too. However, the confidence in revealing pain improves with objective pain scale use and experience (2, 25). Accordingly, being a subjective tool, the ability to use the VAS to detect pain could improve with experience. This aspect was not considered in the present study, since the same experienced clinician performed all the assessments. The reliability of the VAS and its responsiveness to pain level variations were not tested, because of the risk of bias associated with the serial assessment carried out by the same individual after the diagnosis, and this represent a limitation of the present study. On the other hand, the VAS scale is fast to use, and this is its main advantage compared to more complete pain scales, such as the composite ones. The average time for a pain assessment using the CANCRS is 3 minutes. In order to limit the number of the parameters included in the scale, sensitivity and specificity were determined for each parameter of the CANCRS and they were used to determine weighting factors. Weighting factors are useful in order to refine the scale by eliminating parameters with low sensibility or specificity for pain detection (8). This improves the overall sensitivity, specificity, and positive and negative predictive values of the scale. Accordingly, parameters with weighting factors equal to 0 should be excluded, therefore the parameter ‘vocalization’ and the parameter ‘respiratory pattern’ should not be considered in the future when assessing rabbits for abdominal pain. However, their inclusion in the scale should be furtherly investigated for diseases other than gastrointestinal stasis syndrome. All the other parameters have weighting factors equal to 2, except for the parameter ‘palpation of the painful area’ (weighting factor = 1). However, no weighting factor resulted to be equal to 3, pointing out that none of the included parameters has both sensitivity and specificity ≥75%. This might be due to the fact that rabbits tend to mask any signs of pain and it is generally accepted that it is more challenging to assess pain in rabbits than in predatory species (such as cats and dogs) (18). According to Benato et al. (2020), veterinary surgeons state to recognize pain through changes in appetite and abnormal body posture. These parameters could certainly be investigated for future inclusion in a composite pain scale, possibly considering patients with conditions different from gastrointestinal stasis syndrome, which implies changes in appetite by definition.

The responsiveness of the CANCRS to meloxicam administration confirms the internal validity of the scale and makes the scale a valid tool for the monitoring of rabbit with gastrointestinal stasis syndrome. Furthermore, the scale could be used in the future for evaluating other analgesic drugs. However, it should be pointed out that studies concerning pharmacology of analgesic drugs in rabbits other than meloxicam are very limited in the literature (26).

In conclusion, the assessment of internal validity by testing and confirming its responsiveness to analgesia administration, the refinement of the scale by determining weighting factors for each parameter, the determination of a cut-off point for the recognition of pain in rabbits with gastrointestinal statis syndrome, together with the previously established construct validity and reliability, make the CANCRS a useful tool for pain evaluation. Further studies should focus on the reliability of the VAS and on the responsiveness of the VAS to changes in pain level. A refined version of the CANCRS could be useful to determine cut-off values for pain level detection for conditions other than gastrointestinal stasis syndrome.

## Supporting information

S1 Dataset.

